# Spinal Cord Microglia Exhibit Impaired Repair Responses to Myelin Damage

**DOI:** 10.64898/2026.02.23.707565

**Authors:** Matthew C. Zupan, Jack M. Petersen, Addison C. Stover, Nishama De Silva Mohotti, Meredith D. Hartley

## Abstract

**Background:** Multiple sclerosis (MS) is a demyelinating disease of the central nervous system (CNS) that affects both the brain and spinal cord, although the brain has historically received greater attention. In the inducible, oligodendrocyte-specific knockout model of *Myrf*, which results in white matter damage to both the brain and spinal cord, our laboratory previously demonstrated that the brain undergoes partial remyelination following white matter damage, whereas the spinal cord fails to do so. We also observed that brain microglia display a much stronger activation than spinal cord microglia in this model. Microglia regulate remyelination by clearing myelin debris, processing resulting lipids, and modulating an inflammation response.

**Results:** Here, to test our hypothesis, we characterized microglial phenotypes during demyelination in both tissues. The brain exhibited greater early microglial recruitment and higher baseline expression of activation and phagocytic markers, suggesting a primed state for responding to damage. In contrast, spinal cord microglia showed delayed phagocytic marker expression, sustained inflammation, and a predominately amoeboid morphology during demyelination.

**Conclusions:** Together, these findings indicate that brain microglia mount a timely and coordinated response to demyelination that supports remyelination, whereas spinal cord microglia adopt a dysfunctional phenotype that may contribute to impaired myelin repair.

## 1. Introduction

Multiple sclerosis (MS) is characterized by immune-mediated myelin loss in the CNS. Although both brain and spinal cord are affected, the spinal cord has been comparatively understudied, partly due to technical challenges in imaging and analysis. Emerging evidence suggests that lesion progression and repair may differ between these regions, though findings remain inconsistent (1). Some postmortem studies report reduced remyelination in spinal cord lesions compared to brain lesions, whereas others report similar levels (1,2). Experimental autoimmune encephalomyelitis (EAE) models have also shown limited spinal cord remyelination, attributed in part to reduced oligodendrocyte lineage cells (3). Overall, regional differences in remyelination remain incompletely understood.

Using the inducible *Plp1*-iCKO-*Myrf* mouse model, our group previously demonstrated distinct demyelination and remyelination dynamics in brain and spinal cord. In this model, Myrf—a transcription factor essential for oligodendrocyte survival and myelin maintenance—is selectively ablated in oligodendrocytes, resulting in progressive demyelination over 12 weeks followed by partial remyelination in the brain (4–7). Histological and ultrastructural analyses revealed significant myelin loss in both regions during the demyelination phase (7,8). However, while the brain exhibited subsequent remyelination after week 12, the spinal cord showed continued myelin decline without evidence of repair up to week 30 (8).

Lipidomic analysis further highlighted regional differences. Brain cholesterol ester levels increased ∼30-fold at peak demyelination and declined to near normal levels during remyelination. In contrast, spinal cord cholesterol esters increased ∼130-fold and remained elevated, consistent with sustained myelin breakdown (8). Because myelin is cholesterol-rich, demyelination results in excess free cholesterol that becomes esterified and stored in microglia as lipid droplets (9–11). These findings suggested that cholesterol ester dynamics closely track with myelin integrity and pointed toward potential differences in microglial lipid metabolism between the brain and spinal cord.

Microglia, the resident immune cells of the CNS, play essential roles in development, homeostasis, and disease (12,13). During demyelination, activated microglia release pro-inflammatory mediators but are also required for debris clearance and remyelination (14–18). Microglia are highly plastic and respond dynamically to environmental cues, making them attractive but complex therapeutic targets. A clearer understanding of region-specific microglial responses may therefore inform strategies to enhance remyelination (19,20).

In this study, we characterized microglial phenotypes in brain and spinal cord of *Plp1*-iCKO*-Myrf* mice at 6 and 18 weeks post-demyelination, corresponding to active demyelination and remyelination in the brain. Although remyelination is limited in the spinal cord, these timepoints enabled a comparative analysis of microglial responses between the two tissues. Using flow cytometry, immunofluorescence, and morphological analyses, we demonstrated that brain microglia mount an early, coordinated repair response, whereas spinal cord microglia exhibit delayed phagocytic activation, sustained inflammation, and a pathological morphology. Together, these findings demonstrate that in the *Plp1*-iCKO-*Myrf* model with widespread myelin damage, microglia in the brain are better equipped to promote remyelination, whereas spinal cord microglia exhibit dysfunctional responses that may contribute to worse recovery outcomes.

## 2. Materials and Methods

### Animal husbandry

Male and female C57BL/6J *Plp1*-iCKO-*Myrf* mice (*Myrf fl/fl*; *Plp1*-CreERT) (5) were generated by crossing Myrf fl/fl mice (4) with PLP-CreERT mice (21). Animals were housed under standard conditions (24 ± 1 °C, 12-hour light/dark cycle) with food and water ad libitum. Mice were genotyped to assess whether they express the *cre recombinase* transgene (*Cre* positive) or whether they do not express the *cre recombinase* transgene (*Cre* negative) following established protocols (4,21). Both sexes were included. All procedures were approved by the University of Kansas Institutional Animal Care and Use Committee.

### Induction of demyelination

Tamoxifen (T5648, Millipore Sigma) (20 mg/mL in corn oil) was administered intraperitoneally (100 mg/kg) for five consecutive days beginning at 8 weeks of age. Mice were weighed every week and assigned a clinical score adapted from previous studies (5) (1: limp tail or gait tremor; 2: partial hind limb weakness; 3: hind limb paralysis (could not hang from hind limbs); and 4: ataxia and severe hindlimb paralysis). Animals were euthanized at week 6 or 18 post-tamoxifen.

### Flow cytometry

For flow cytometry analysis, mice were anesthetized with an 80 mg/mL and 12 mg/mL ketamine/xylazine solution and then perfused transcardially with 40 mL of ice-cold HBSS without Ca^2+^ and Mg^2+^. Brain and spinal cord tissues were collected from the mice, homogenized in a Dounce homogenizer, and a murine leukocyte population containing microglia was isolated using a Percoll gradient as described (22). The resulting cells were incubated in an FC block solution (Tonbo 70-0161, 1:100) for 10 minutes and then stained in an antibody mix for 30 minutes which consisted of CD45 violetFluor 450 (Tonbo 75-0451, 1:75), CD11b PerCP-Cyanine5.5 (Tonbo 65-0112, 1:75), Brilliant Violet 605 anti-mouse CD16 (Biolegend 158027, 1:75), FITC anti-mouse CD68 (Biolegend 137006, 1:75), APC/Fire 810 anti-mouse CX3CR1 (Biolegend 149054, 1:75), APC anti-mouse F4/80 (Biolegend 123116, 1:75), APC/Fire 750 anti-mouse CD73 (Biolegend 127221, 1:75), PE anti-mouse TREM2 (R&D Systems FAB17291P, 1:75), and viability dye (Tonbo 13-0871, 1:1000). Single stain controls were made using compensation beads (Biolegend, 424602) and following the kit instructions. The beads, brain and spinal cord samples were then run on a flow cytometer (Cytek Aurora full spectrum flow cytometer). The unstained cells and single stain controls were first run to set the positive controls for each antibody, and then 50,000 cells were run for each sample.

### Immunofluorescence

For immunofluorescence imaging, mice were anesthetized with an 80 mg/mL and 12 mg/mL ketamine/xylazine solution and then perfused transcardially with 10 mL HBSS without Ca^2+^ and Mg^2+^ followed by 40 mL of 4% PFA in HBBS. Brains and spinal cords were stored in 4% PFA at 4 ° C. Free-floating sections (40 µm) were obtained using a Vibratome. The sections were permeabilized in HBSS/0.01% sucrose + 0.1% saponin (HBSS/Su/Sap) + 0.05% Triton X-100 for 30 minutes at room temperature followed by a 10-minute rinse in HBSS. The sections were then incubated in blocking buffer consisting of HBSS/Su/Sap + 3% normal donkey serum for 2 hours at room temperature followed by an incubation with primary antibodies diluted in blocking buffer overnight at 4 ° C. The next day, the tissue sections were rinsed three times for 10 minutes with HBSS and then incubated in a solution of secondary antibodies diluted in HBSS/Su/Sap for 3.5 hours and rinsed three times for 10 minutes in HBSS. They were then incubated in a solution of DAPI (0.5 µg/mL) dissolved in HBSS for 1 hour. The tissue sections were finally rinsed two times for 10 minutes with HBSS and then mounted onto subbed slides using ProLong Diamond antifade mounting media (Life Technologies P36961).

The following primary antibodies were used: mouse anti-MerTK (Santa Cruz Biotechnology sc-365499, 1:50), sheep anti-Trem2 (R&D System AF1729, 1:50), rabbit anti-IBA1 (Wako 019-19741, 1:500), rat anti-CD68 (Abcam ab53444, 1:50), mouse anti-iNOS (BD Biosciences 610329, 1:100), and rabbit anti-liver arginase (Abcam ab91279, 1:100). The following secondaries were used: donkey anti-sheep 488 Alexa Fluor (Invitrogen A-11015, 1:200), donkey anti-mouse 647 Alexa Fluor (Invitrogen A-31571, 1:200), donkey anti-rabbit 555 Alexa Fluor (Invitrogen A-31572, 1:200), and donkey anti-rat 488 Alexa Fluor (Invitrogen A-48269, 1:200).

Brains and spinal cords stained for immunofluorescence were imaged using a Nikon 2037 CSU-W1 spinning disk confocal microscope. The medial corpus callosum and the ventral and dorsal regions of the lumbar spinal cord were imaged with a 20x/NA 0.8 objective (709.81 x 744.04 microns, 2198 x 2304 pixels). For each section, a Z-stack with a step size of 4 µm was acquired. For the colocalization studies (MerTK/TREM2-IBA1 and iNOS/Arg1-CD68), sections were imaged with a 4xSoRa objective and Z-stacks with a step size of 0.8 µm were obtained. The images were analyzed using Image J, where the Z-stacks were compressed into a single maximum intensity projection. The mean intensities of the staining and the staining area at a specific optimized threshold were recorded. The same threshold was used for all images analyzed in a single batch for each marker. A colocalization analysis was performed using thresholding to determine the percent area of IBA1+ cells that overlapped with MerTK/TREM2 staining and the percent area of CD68+ cells that overlapped with iNOS/Arg1 staining.

### Microglial morphology

IBA1-stained microglia were analyzed using the MicrogliaMorphology ImageJ macro and MicrogliaMorphologyR package to quantify 27 morphological features per cell. Clustering analysis identified distinct morphological classes. Instructions were followed per the provided tutorial and the supporting code can be found on GitHub at https://github.com/ciernialab/MicrogliaMorphology (23). The data obtained by the MicrogliaMorphology ImageJ macro was analyzed and visualized using the complimentary R package developed by the Ciernia lab, MicrogliaMorphologyR. The source code for MicrogliaMorphologyR as well as instructions on how to use the code can be found on GitHub at https://github.com/ciernialab/MicrogliaMorphologyR (23).

### Statistics

Group sizes were based on prior studies and preliminary data. Animals were randomly assigned with balanced sex distribution when possible. Two- or three-way ANOVA with Tukey’s post hoc tests were used as appropriate. P ≤ 0.05 was considered significant.

## 3. Results

### Brain microglia are recruited earlier and more robustly during disease than spinal cord microglia

Brain and spinal cord tissue from *Plp1*-iCKO-*Myrf* mice at time points of week 6 and week 18 post-tamoxifen were immunofluorescently stained for IBA-1, a macrophage marker (Fig. 1A, B). The IBA-1 signal was significantly greater in the brain than in the spinal cord at week 6, indicating an earlier response by microglia to demyelination. By week 18, the spinal cord exhibited higher IBA-1 levels, consistent with delayed and prolonged microglial accumulation (Fig. 1 C).

**Figure 1.**
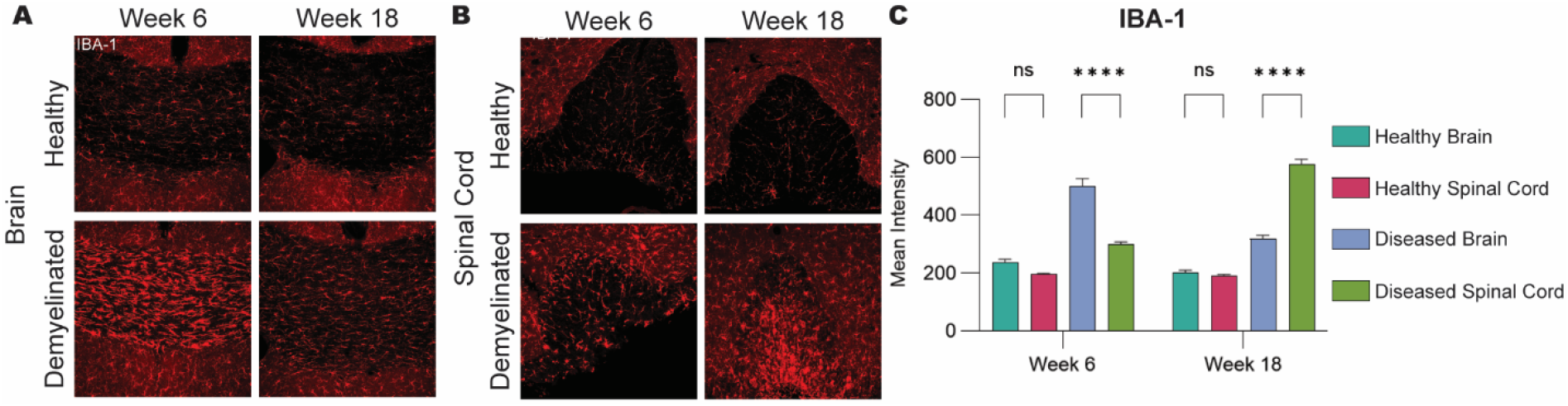
IBA-1 is differentially expressed in the brain and spinal cord during myelin damage. A) Representative immunofluorescence images of the medial corpus callosum in the brain stained with antibodies for IBA-1. (B) Representative immunofluorescence images of the dorsal region in the lumbar spinal cord stained with antibodies for IBA-1. (C) IBA-1 mean intensities graphed to depict the comparison between brain and spinal cord. Mean intensities were obtained using Fiji Image J software. Error bars represent standard error of the mean. Statistical analysis was performed using a 3-way ANOVA with Tukey’s post-hoc analysis, n = 5 – 15 where each sample represents a different animal, * p < 0.05, ** p < 0.01, *** p < 0.001, **** p <0.0001.

Next, microglia were isolated from the brain and spinal cord of *Plp1*-iCKO-*Myrf* mice at week 6 and week 18 post-tamoxifen using a Percoll gradient and flow cytometry (22). The flow cytometry analysis was performed by gating the total leukocyte population was gated away from any debris, then single cells were gated away from any multiplets. Next, single cells were gated by viability and for their expression level of surface marker CD45. Cells expressing high levels of CD45 (CD45 high) are classified as infiltrating macrophages, whereas microglia are classified as having low levels of CD45 (CD45 low). Lastly, the CD45 low population was gated to include cells positive for expression of surface marker CD11b (Fig. 2A-D). Cells that are CD45 low and CD11b+ are generally accepted to be a pure microglia population (22,24). We then determined the percentage of cells that are microglia in relation to all the single cells in the sample to get a measure of the microglia population in each condition. We directly compared the brain and spinal cord microglia percentages and observed that at week 6 during disease, the brain has a significantly higher percentage of microglia cells than the spinal cord (Fig. 2E). Previous studies have supported that microglia density is positively associated with remyelination and OPC levels (18,25). These data suggest that rapid microglial recruitment in the brain may support efficient debris clearance and remyelination, whereas delayed recruitment in the spinal cord may contribute to impaired repair.

**Figure 2.**
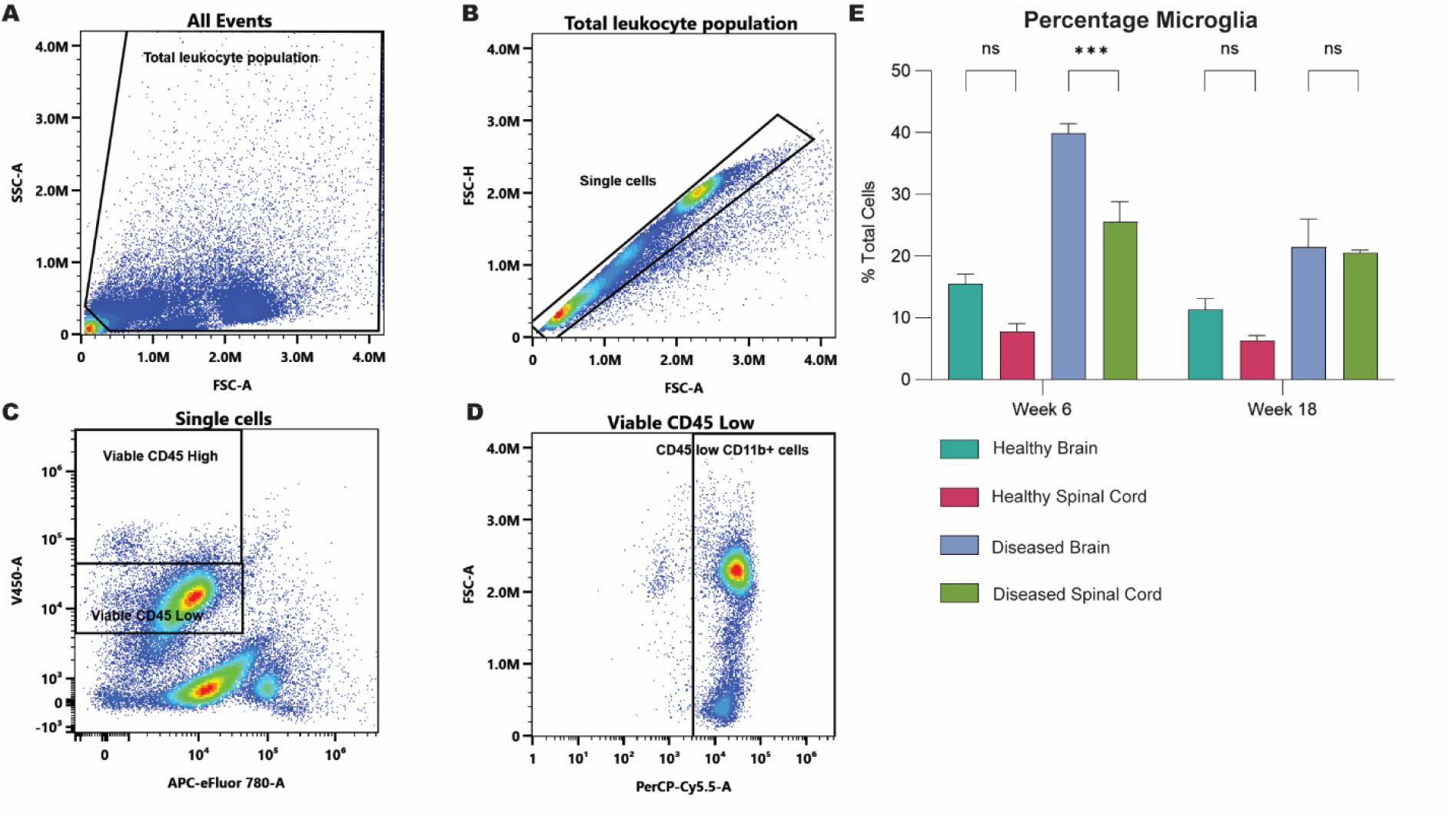
Flow cytometry analysis reveals that microglia population numbers vary across brain and spinal cord during myelin damage. (A). Flow cytometry workflow. The total leukocyte population was gated away from debris. (B) Single cells were then gated away from any multiplets. (C) Viable cells expressing low levels of CD45 were gated away from other cells and infiltrating macrophages. (D) Lastly, cells expressing low levels of CD45 and positive for CD11b were selected for the analysis, as they are microglia. (E) Microglia as a percentage of the total single cells in the sample graphed to highlight the comparison between brain and spinal cord. Data represents mean and error bars represent standard error of the mean. Statistical analysis was performed using a 3-way ANOVA with Tukey’s post-hoc analysis, n = 7 – 10 where each sample represents a different animal, * p < 0.05, ** p < 0.01, *** p < 0.001, **** p <0.0001.

### Microglia in the diseased spinal cord adopt a pathological amoeboid morphology

Microglia are highly heterogeneous and dynamic cells that display diverse morphological and functional states depending on the signals they detect from the CNS environment (26). Historically, microglia with a ramified morphology have been classified as the “resting state” microglia and reactive microglia with retracted processes and larger somas have been classified as “activated” microglia that are associated with inflammation (26,27). Another morphology that microglia can adopt is an “ameboid” one, characterized by a round body with no processes (27). Microglia exhibiting this morphology are generally associated with more severe pathology.

To investigate differences in microglia morphology between the brain and the spinal cord over the course of demyelination, IBA-1 immunofluorescence images acquired from tissues slices were analyzed using a microglia morphology analysis toolset developed by the Ciernia lab (Fig. 3A) (23). Using this toolset, four distinct microglia clusters were identified that could be differentiated based on their unique morphology features (Fig. 3B). Cluster 1 was labeled as ameboid due to its high circularity and smallest branch lengths. Cluster 2 was labeled as ramified due to its long but sparsely branched processes. Cluster 3 was labeled as hyper-ramified due to its high branching complexity with thin processes. Cluster 4 was labeled as rod-like due to elongated and highly bipolar morphology.

**Figure 3.**
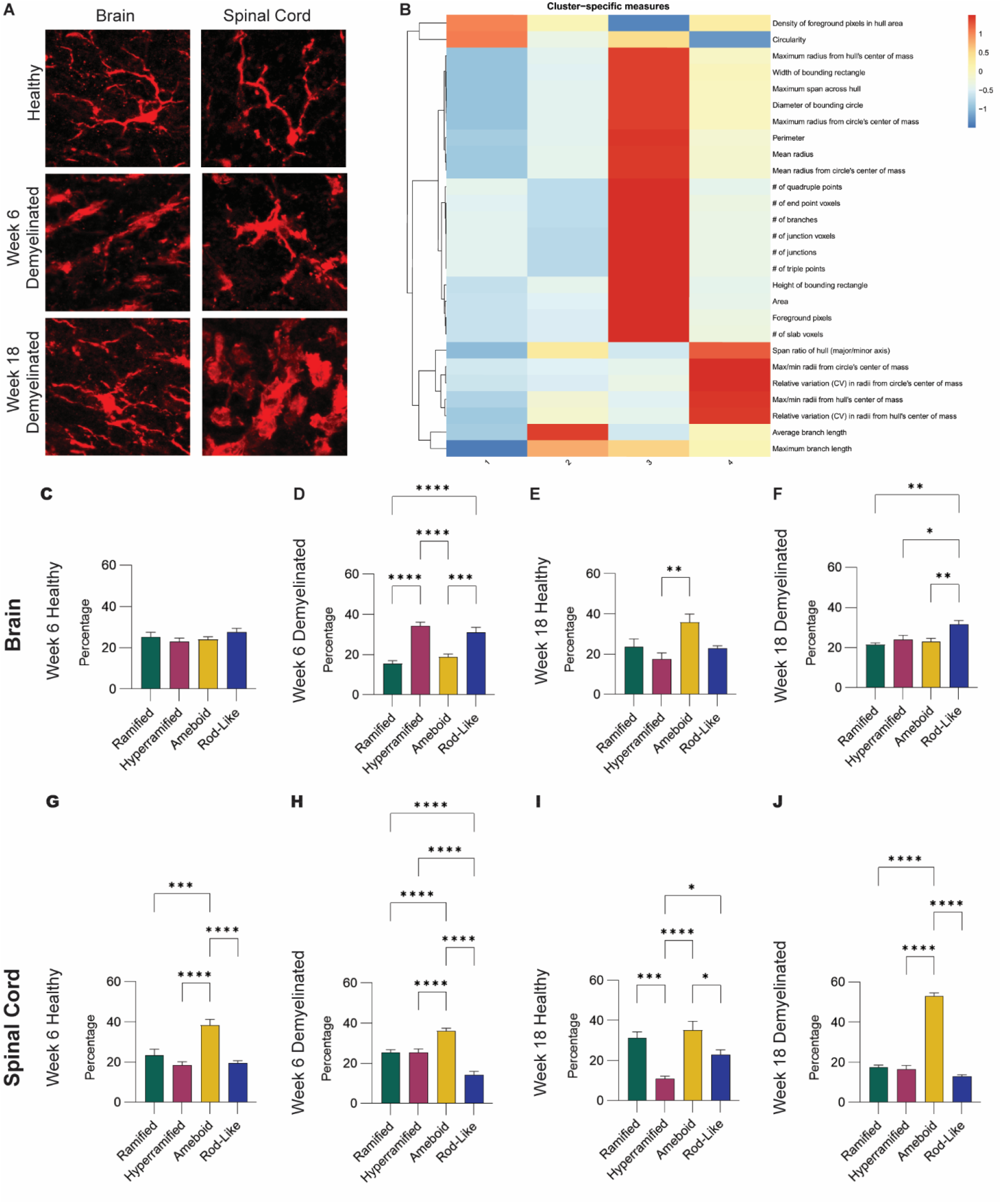
Microglia morphology is highly dynamic across the brain and spinal cord during myelin damage. (A). Representative images of microglia morphology in brain and spinal cord across the time course of demyelination. (B) Comparison of microglia morphology features across clusters. Cluster 1 = ameboid, Cluster 2 = ramified, Cluster 3 = hyper-ramified, and Cluster 4 = rod-like. (C-J) Percentage of microglia exhibiting a specific morphology in the brain and spinal cord across all healthy and diseased conditions. Data represents mean and error bars represent standard error of the mean. Statistical analysis was performed using a one-way ANOVA with Tukey’s post-hoc analysis, n = 7 – 10 where each sample represents a different animal, * p < 0.05, ** p < 0.01, *** p < 0.001, **** p <0.0001.

During active demyelination at week 6, brain microglia predominantly exhibited hyper-ramified and rod-like morphologies and by week 18 they revert to a ramified morphology, with some still exhibiting a rod-like morphology (Fig 3. D, F). A hyper-ramified morphology is considered an intermediate transitionary state between resting and reactive states and rod-like is considered a pathological state of reactive microglia (28,29). So overall, these data demonstrate that over the course of disease in the brain, microglia transition towards a reactive state during demyelination and begin to return to a resting state during active remyelination.

In contrast, spinal cord microglia were enriched in the ameboid cluster, particularly at week 18 after demyelination (Fig. 3G-J). An ameboid morphology is commonly associated with chronic pathology and inflammatory activation (30,31). Although morphology alone does not define function, the predominance of amoeboid microglia in the spinal cord suggests a maladaptive response during sustained myelin damage.

### Brain microglia exhibit higher baseline activation marker expression

To further characterize microglia function and behavior in the brain and spinal cord, flow cytometry was employed to investigate the expression of six surface proteins that play important roles during neurological disease (Table 1).

**Table 1.**
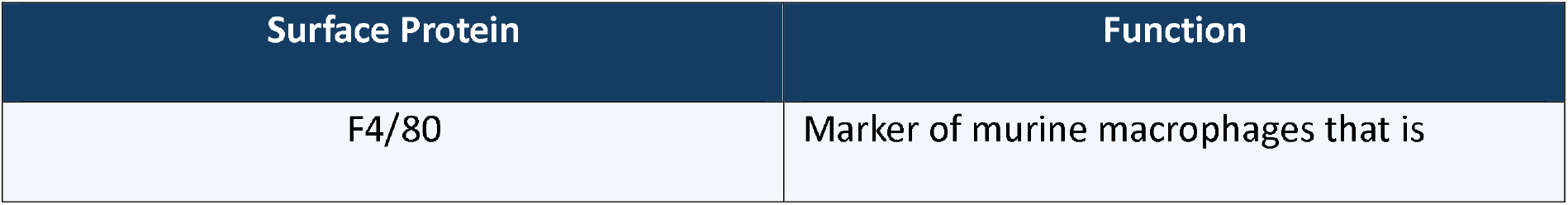
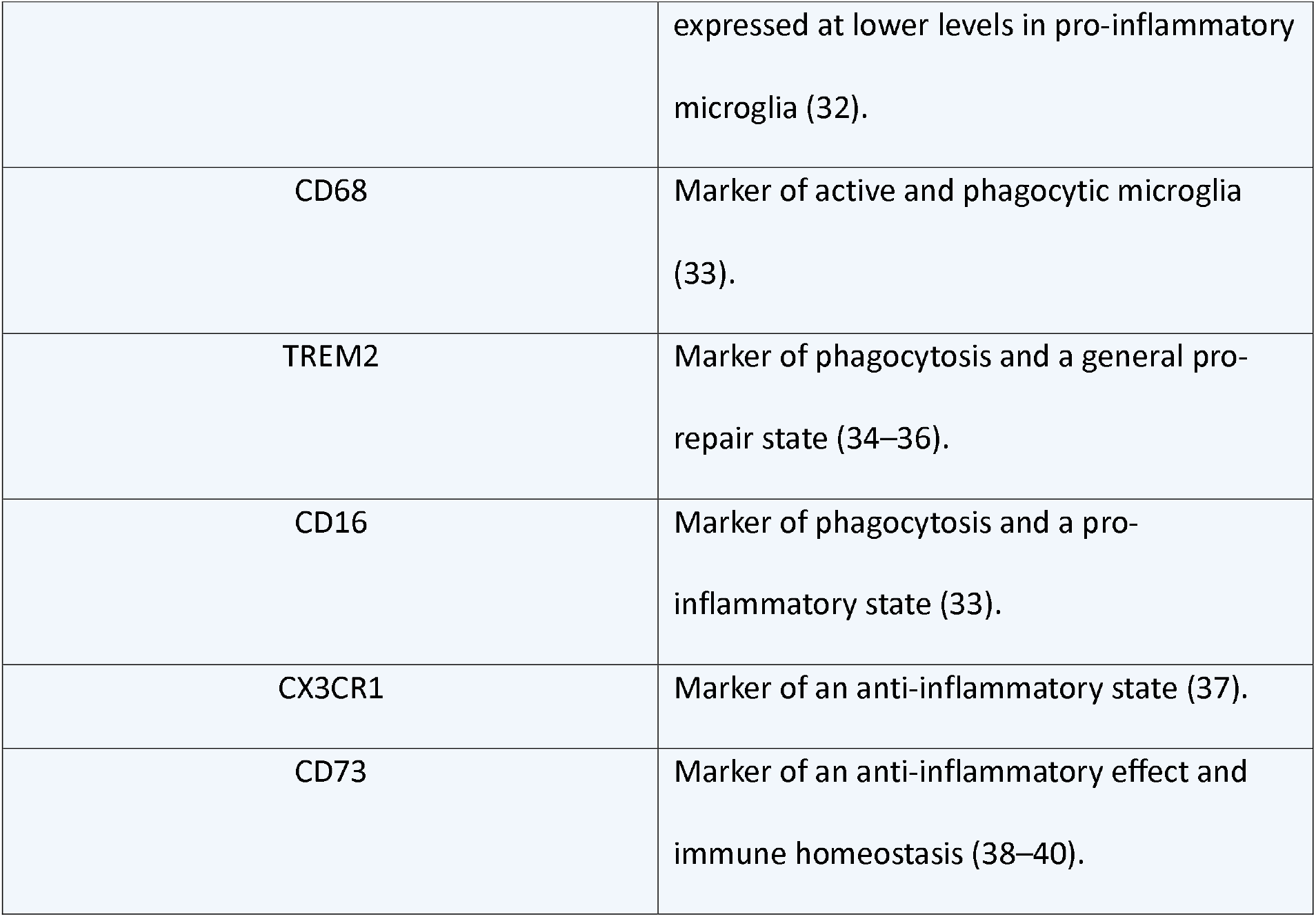
Surface proteins selected in the flow cytometry marker panel to investigate microglia phenotype.

| Surface Protein | Function |
| --- | --- |
| F4/80 | Marker of murine macrophages that is |
|  | expressed at lower levels in pro-inflammatory microglia (32). |
| CD68 | Marker of active and phagocytic microglia (33). |
| TREM2 | Marker of phagocytosis and a general pro-repair state (34–36). |
| CD16 | Marker of phagocytosis and a pro-inflammatory state (33). |
| CX3CR1 | Marker of an anti-inflammatory state (37). |
| CD73 | Marker of an anti-inflammatory effect and immune homeostasis (38–40). |

Flow cytometry demonstrated that healthy brain microglia expressed higher levels of all six analyzed markers compared healthy to spinal cord microglia, suggesting a primed baseline state (Fig. 4). At week 6 during disease, marker expression was similar between brain and spinal cord microglia, except for CD73, which was more highly expressed in brain microglia. At week 18 during disease, all markers were more highly expressed in brain microglia compared to spinal cord microglia, except for F4/80 and CX3CR1, which both showed similar expression levels between the two tissues. Notably, CD73 expression was consistently elevated in brain microglia across all conditions. Given the role of CD73 in regulating inflammatory balance, its reduced expression in the spinal cord may predispose this region to excessive inflammation.

**Figure 4.**
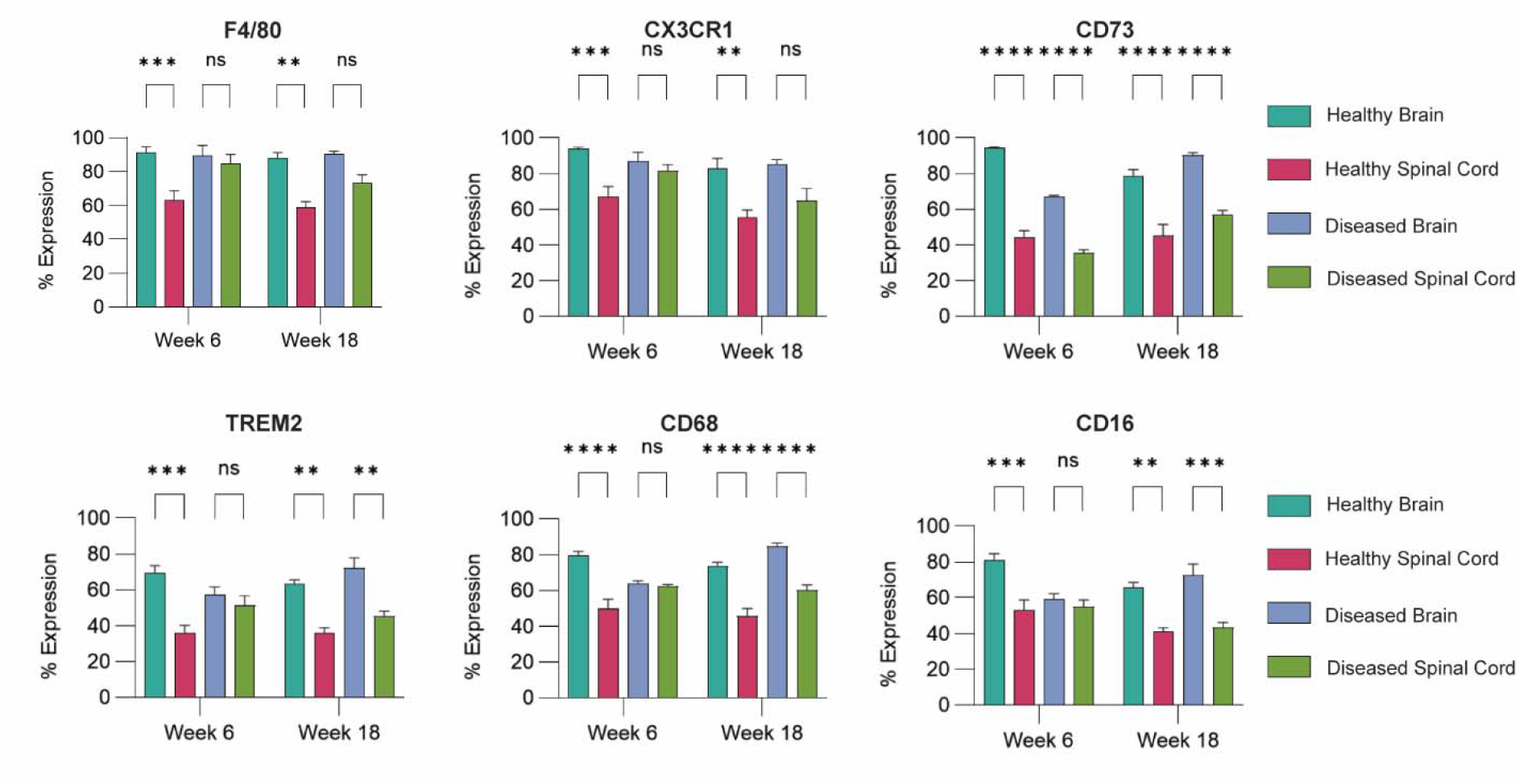
Expression of selected surface protein markers is generally higher in brain microglia compared to spinal cord microglia. Expression of each marker as a percentage of the total microglia in the sample. Data represents mean and error bars represent standard error of the mean. Statistical analysis was performed using a 3-way ANOVA with Tukey’s post-hoc analysis, n = 3 – 10 where each sample represents a different animal, * p < 0.05, ** p < 0.01, *** p < 0.001, **** p <0.0001.

Overall, the flow cytometry analysis demonstrated that brain microglia generally had a higher baseline expression level of these six important proteins under healthy conditions (Fig. 4). Higher expression of these markers could prime the brain microglia to be better prepared to handle myelin damage compared to spinal cord microglia.

### Brain microglia mount an early phagocytic response, whereas the spinal cord response is delayed

To further elucidate this tissue-based microglial response to demyelination, immunofluorescence staining was performed on brain and spinal cord tissue sections of *Plp1*-iCKO-*Myrf* mice at week 6 and week 18 post-tamoxifen (Fig. 5 A, C). We first stained for two proteins that are involved in phagocytosis, MerTK and TREM2. MerTK plays an essential role in microglial phagocytosis and migration (41). It binds and is activated by phosphatidylserine, which is abundantly found in myelin, and subsequently activates debris engulfment (42,43). MerTK knockout experiments have led to chronic inflammation and deficiencies in microglia phagocytosis and oligodendrocyte differentiation (41,42). TREM2 is a microglia transmembrane protein that acts as a lipid sensor and promotes microglia activation in response to myelin damage by activating a kinase signaling cascade that induces the transcription of genes involved in phagocytosis, oligodendrocyte maturation and inflammation regulation (35,44,45).

**Figure 5.**
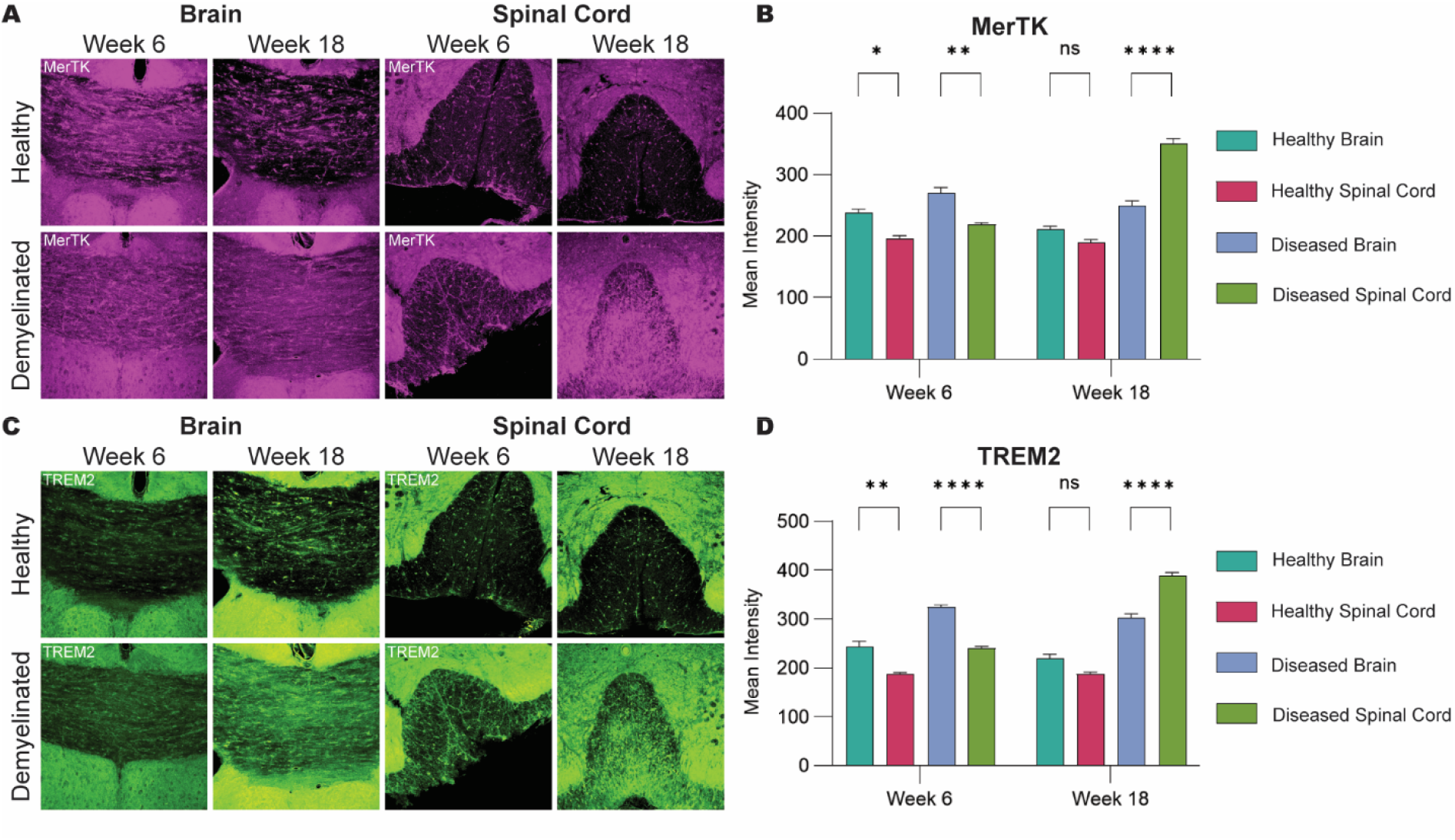
Phagocytosis markers show an earlier response in the demyelinating brain compared to the spinal cord. (A, C). Representative immunofluorescence images of the medial corpus callosum in the brain and the dorsal region in the lumbar spinal cord stained with antibodies for MerTK and TREM2. (B, D) Mean intensities graphed to depict the comparison between brain and spinal cord. Mean intensities were obtained using Fiji Image J software. Error bars represent standard error of the mean. Statistical analysis was performed using a 3-way ANOVA with Tukey’s post-hoc analysis, n = 5 – 15 where each sample represents a different animal, * p < 0.05, ** p < 0.01, *** p < 0.001, **** p <0.0001.

MerTK and TREM2 immunostaining revealed elevated expression in the brain during demyelination, consistent with active debris clearance. In contrast, spinal cord expression of these markers showed no changes at week 6 and was only increased at week 18, after prolonged myelin damage (Fig. 5 B, D). These findings indicate that the brain initiates phagocytic mechanisms early, while the spinal cord exhibits delayed activation.

### Spinal cord microglia exhibit a sustained pro-inflammatory response to demyelination

To assess the inflammation response of the brain and spinal cord to demyelination, tissue sections were immunofluorescently stained for inducible nitric oxide synthase (iNOS) and arginase-1 (Arg1) (Fig. 6A). iNOS produces nitric oxide which ultimately creates an inflammatory environment (46). There is evidence that some level of iNOS is necessary during demyelination to regulate inflammation and autoimmunity (47). However, excessive nitric oxide production is neurotoxic (46). In response to this overproduction of nitric oxide, macrophages upregulate arginases, which compete for the same substrate as iNOS, arginine, thereby limiting iNOS function and promoting an anti-inflammatory environment (46). This switch from strong expression of iNOS, commonly observed at the onset of demyelination, to Arg1 expression is considered to be critical to initiate remyelination, resolve neuroinflammation, and restore tissue integrity (46). The ratio of iNOS and Arg1 expression can indicate whether the environment is leaning towards a more pro-inflammatory or anti-inflammatory phenotype.

**Figure 6.**
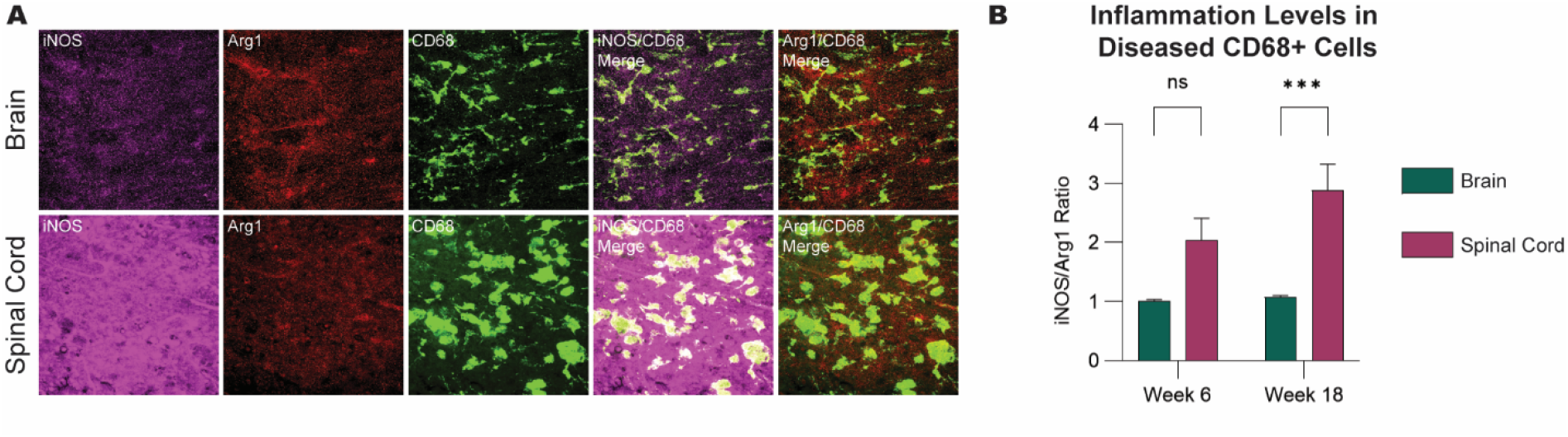
Co-localization analysis of active microglia reveal that spinal cord microglia show an extended inflammatory response during demyelination. (A) Representative brain and spinal cord immunofluorescence images used for the co-localization analysis of CD68+ cells with iNOS expression and Arg1 expression, respectively. (B) Ratio of percent colocalization of CD68+ cells with iNOS to percent colocalization of CD68+ with Arg1. Data represents mean and error bars represent standard error of the mean. The co-localization analysis was performed using Fiji Image J software. Statistical analysis was performed using a 2-way ANOVA with Tukey’s post-hoc analysis, n = 9 where 3 biological samples were analyzed in triplicate, * p < 0.05, ** p < 0.01, *** p < 0.001, **** p <0.0001.

Colocalization analysis of iNOS and Arg1 within CD68+ microglia revealed a marked shift toward iNOS dominance in the diseased spinal cord at week 18. The iNOS:Arg1 ratio increased approximately threefold, indicating sustained pro-inflammatory response in spinal cord microglia. In contrast, brain microglia showed a more balanced profile consistent with resolution of inflammation (Fig. 6B).

## 4. Discussion

Efficient myelin repair requires timely debris clearance, as myelin debris contains inhibitory factors that prevent OPC recruitment and differentiation (48). Activated microglia have been shown to be essential for OPC migration, proliferation, and differentiation, likely through the secretion of neurotrophic factors (49,50). In this study, we examined a panel of microglia markers to assess the phenotypes of brain and spinal cord microglia during demyelination (Table 1). Looking back in more detail at some of the markers depicted in Table 1, CX3CR1 has been shown to play a role in microglia migration as well as myelin debris clearance (51,52), TREM2 has been shown to promote myelin debris clearance and remyelination (48), CD68 is considered a marker of actively phagocytic microglia (33), and CD16 expressing microglia release pro-inflammatory cytokines that are important for initial microglia activation and recruitment (53,54). All these markers are involved in responding to myelin damage, and we observed that healthy brain microglia have a higher baseline expression of these markers compared to healthy spinal cord microglia (Fig. 4) suggesting that brain microglia are better equipped to respond to myelin injury.

The expression of CD73 from the flow cytometry data was particularly interesting, seeing as CD73 expression in the brain microglia was considerably higher than in the spinal cord microglia across all conditions, both in control and demyelinated mice (Fig. 4). CD73 plays an important role in maintaining immune homeostasis by balancing pro-inflammatory and immunosuppressive signals (38–40). Overall, the higher expression of CD73 in brain microglia suggests that the brain is better equipped to manage inflammation during disease, and that the spinal cord may be more susceptible to inflammation due to reduced CD73. This further supports that the brain microglia are better equipped to lead an efficient response to demyelination compared to spinal cord microglia.

Additionally, the quick response to the onset of demyelination in the brain is highlighted by an increase in microglia recruitment (Fig. 1, 2) and an increase in MerTK and TREM2 expression (Fig 5.) TREM2 has extensively been shown to promote myelin debris clearance and remyelination (36,55–57) and MerTK has been shown to play a role in microglia recruitment and phagocytosis during demyelination (41). Therefore, the early increase in TREM2 and MerTK expression at week 6 in the diseased brain implies that the brain microglia respond more quickly to myelin damage by phagocytosing the debris leading to more efficient remyelination. In contrast, the diseased spinal cord does not exhibit a strong microglia recruitment or increased TREM2 and MerTK expression at the onset of demyelination (Fig. 1,2), meaning that the spinal cord microglia are not effectively responding to myelin damage and phagocytosing debris, which is consistent with the lack of efficient remyelination observed in the demyelinated spinal cord. This phenotype is further represented by the strong pro-inflammatory shift of the diseased spinal cord at week 18 (Fig. 6).

Furthermore, the increasingly ameboid morphology and pro-inflammation shift of microglia in the diseased week 18 spinal cord supports the lack of appropriate early response to demyelination and shows a potentially defective phenotype of spinal cord microglia. An ameboid microglia morphology is typically found in chronically inflamed pathological tissue and is associated with advanced stages of neurological disease. Additionally, ameboid microglia can release neurotoxic cytokines, leading to a strongly pro-inflammatory and pathological environment (30). Since ameboid microglia are the most prominent microglia morphology in the diseased spinal cord at both timepoints, we can conclude that the diseased spinal cord is continually advancing towards a neurotoxic and pathological state. In contrast, the diseased brain displays a larger proportion of rod-like microglia, which are active phagocytic microglia associated with earlier stages of disease (30). The brain microglia are able to effectively handle demyelination without ever reaching a severe pathological ameboid state.

These differences between brain and spinal cord microglia can be attributed to a few possible causes. Developmental myelination ends earlier in the spinal cord than in the brain (58), so it is possible that spinal cord microglia then enter a quiescent state as there is less for them to do. This could explain the protein expression differences in brain and spinal cord microglia even in healthy tissue. In this quiescent state, they could be less prepared to respond to the acute damage that occurs during demyelination. Additionally, the spinal cord contains more heavily myelinated axons compared to the brain (59). Therefore, demyelination causes a larger amount of myelin debris to accumulate which could overwhelm the spinal cord microglia, causing them to become lipid-laden and dysfunctional (60,61).

In conclusion, our data demonstrate region-specific microglial responses to demyelination. Brain microglia exhibit rapid recruitment, early phagocytic activation, and subsequent resolution— features consistent with effective remyelination. In contrast, spinal cord microglia display delayed phagocytic activation, sustained inflammation, and predominately amoeboid morphology, coinciding with failed remyelination. Effective remyelination requires that microglia transition from an early, predominantly pro-inflammatory signature to a later, pro-repair state (62). In the diseased spinal cord at week 18, the persistence of amoeboid morphology, elevated expression of MerTK and TREM2, and increased iNOS levels all point to a maladaptive, neurotoxic microglial profile rather than a reparative one (30,31,46,63,64). Taken together, these findings suggest that spinal cord microglia become functionally impaired during chronic white matter damage in this model, failing to execute the phenotypic shift necessary for remyelination.

These findings raise the possibility that intrinsic differences between brain and spinal cord microglia—or region-specific environmental cues—shape repair capacity. Targeting microglial phenotype modulation may therefore represent a promising therapeutic strategy to enhance spinal cord remyelination.

## Abbreviations

CNS: central nervous system
ACAT1: acetyl-CoA-acyltransferase1
OPC: oligodendrocyte precursor cell
EM: electron microscopy
TREM2: triggering receptor-expressed on myeloid cells 2
MerTK: MER proto-oncogene tyrosine kinase
iNOS: inducible nitric oxide synthase
ARG-1: arginase-1

## Acknowledgments

Immunofluorescence images were acquired at the Kansas Intellectual and Developmental Disabilities Research Center imaging core facility using the Nikon CSU-W1 spinning disk confocal microscope that was funded by NIH S10 OD 032207 at the University of Kansas Medical Center.

Flow cytometry was performed at the University of Kansas Flow Cytometry Core with guidance and assistance from Dr. Robin Orozco and Dr. Peter McDonald.

Technical assistance was provided by Jenna Williams, Rashmi Binjawadagi, and Hiroko Kobayashi.

## Author Information

### Author Affiliations

**University of Kansas, Department of Chemistry, Lawrence, KS 66044, USA**.

Matthew C. Zupan, Nishama D.S. Mohotti, Meredith D. Hartley

### Contributions

All experiments and analyses were performed by M.Z. J.MP and A.C.S. provided assistance with transcardiac perfusions and tissue homogenization. Tissue sections for immunofluorescence analysis were prepared by N.D.S.M. M.Z. and M.D.H. conceived the project, designed the experiments, and wrote the manuscript.

## Ethics Declaration

### Ethics approval and consent to participate

All animal experiments were approved by IACUC committees at the University of Kansas.

### Consent for publication

Not applicable.

## References

1. Kreiter D, Postma AA, Hupperts R, Gerlach O. Hallmarks of spinal cord pathology in multiple sclerosis. Journal of the Neurological Sciences. 2024 Jan;456:122846. doi:10.1016/j.jns.2023.122846

2. Frischer JM, Weigand SD, Guo Y, Kale N, Parisi JE, Pirko I, et al. Clinical and pathological insights into the dynamic nature of the white matter multiple sclerosis plaque. Annals of Neurology. 2015 Nov;78(5):710–21. doi:10.1002/ana.24497

3. Hampton DW, Anderson J, Pryce G, Irvine KA, Giovannoni G, Fawcett JW, et al. An experimental model of secondary progressive multiple sclerosis that shows regional variation in gliosis, remyelination, axonal and neuronal loss. Journal of Neuroimmunology. 2008 Sep;201–202:200–11. doi:10.1016/j.jneuroim.2008.05.034

4. Emery B, Agalliu D, Cahoy JD, Watkins TA, Dugas JC, Mulinyawe SB, et al. Myelin Gene Regulatory Factor Is a Critical Transcriptional Regulator Required for CNS Myelination. Cell. 2009 Jul;138(1):172–85. doi:10.1016/j.cell.2009.04.031

5. Koenning M, Jackson S, Hay CM, Faux C, Kilpatrick TJ, Willingham M, et al. Myelin Gene Regulatory Factor Is Required for Maintenance of Myelin and Mature Oligodendrocyte Identity in the Adult CNS. J Neurosci. 2012 Sep 5;32(36):12528–42. doi:10.1523/JNEUROSCI.1069-12.2012

6. Doerflinger NH, Macklin WB, Popko B. Inducible site-specific recombination in myelinating cells. Genesis. 2003 Jan;35(1):63–72. doi:10.1002/gene.10154

7. Hartley MD, Banerji T, Tagge IJ, Kirkemo LL, Chaudhary P, Calkins E, et al. Myelin repair stimulated by CNS-selective thyroid hormone action. JCI Insight. 2019;4(8). doi:10.1172/JCI.INSIGHT.126329

8. De Silva Mohotti N, Williams JM, Binjawadagi R, Kobayashi H, Dedunupitiya D, Petersen JM, et al. Temporal dynamics of CNS cholesterol esters correlate with demyelination and remyelination [Internet]. Neuroscience; 2025 [cited 2025 Oct 1]. Available from: http://biorxiv.org/lookup/doi/10.1101/2025.09.06.674663 doi:10.1101/2025.09.06.674663

9. Wender M, Filipek-Wender H, Stanisławska J. Cholesteryl esters of the brain in demyelinating diseases. Clinica Chimica Acta. 1974 Aug;54(3):269–75. doi:10.1016/0009-8981(74)90245-9

10. Fewster ME, Mead JF, Wolfgram FJ, Tourtellotte WW. Cholesterol Esters in Myelin Isolated from Cerebral White Matter of Patients with Multiple Sclerosis. Experimental Biology and Medicine. 1970 Mar 1;133(3):795–800. doi:10.3181/00379727-133-34566

11. Ralhan I, Chang CL, Lippincott-Schwartz J, Ioannou MS. Lipid droplets in the nervous system. Journal of Cell Biology. 2021 Jul 5;220(7):e202102136. doi:10.1083/jcb.202102136

12. Li Q, Barres BA. Microglia and macrophages in brain homeostasis and disease. Nat Rev Immunol. 2018 Apr;18(4):225–42. doi:10.1038/nri.2017.125

13. Thion MS, Ginhoux F, Garel S. Microglia and early brain development: An intimate journey. Science. 2018 Oct 12;362(6411):185–9. doi:10.1126/science.aat0474

14. Luo C, Jian C, Liao Y, Huang Q, Wu Y, Liu X, et al. The role of microglia in multiple sclerosis. NDT. 2017 Jun;Volume 13:1661–7. doi:10.2147/NDT.S140634

15. Olah M, Amor S, Brouwer N, Vinet J, Eggen B, Biber K, et al. Identification of a microglia phenotype supportive of remyelination. Glia. 2012 Feb;60(2):306–21. doi:10.1002/glia.21266

16. Voß EV, Škuljec J, Gudi V, Skripuletz T, Pul R, Trebst C, et al. Characterisation of microglia during de- and remyelination: Can they create a repair promoting environment? Neurobiology of Disease. 2012 Jan;45(1):519–28. doi:10.1016/j.nbd.2011.09.008

17. Wolswijk G. Chronic Stage Multiple Sclerosis Lesions Contain a Relatively Quiescent Population of Oligodendrocyte Precursor Cells. J Neurosci. 1998 Jan 15;18(2):601–9. doi:10.1523/JNEUROSCI.18-02-00601.1998

18. Patani R, Balaratnam M, Vora A, Reynolds R. Remyelination can be extensive in multiple sclerosis despite a long disease course. Neuropathology Appl Neurobio. 2007 Jun;33(3):277–87. doi:10.1111/j.1365-2990.2007.00805.x

19. Lloyd AF, Miron VE. The pro-remyelination properties of microglia in the central nervous system. Nature Reviews Neurology. 2019;15(8). doi:10.1038/s41582-019-0184-2

20. Pons V, Rivest S. Beneficial Roles of Microglia and Growth Factors in MS, a Brief Review. Front Cell Neurosci. 2020 Sep 23;14:284. doi:10.3389/fncel.2020.00284

21. Doerflinger NH, Macklin WB, Popko B. Inducible site-specific recombination in myelinating cells. Genesis. 2003 Jan;35(1):63–72. doi:10.1002/gene.10154

22. Garcia JA, Cardona SM, Cardona AE. Isolation and Analysis of Mouse Microglial Cells. CP in Immunology. 2014 Feb;104(1). doi:10.1002/0471142735.im1435s104

23. Kim J, Pavlidis P, Ciernia AV. Development of a High-Throughput Pipeline to Characterize Microglia Morphological States at a Single-Cell Resolution. eNeuro. 2024 Jul;11(7):ENEURO.0014-24.2024. doi:10.1523/ENEURO.0014-24.2024

24. Brandenburg S, Blank A, Bungert AD, Vajkoczy P. Distinction of Microglia and Macrophages in Glioblastoma: Close Relatives, Different Tasks? IJMS. 2020 Dec 27;22(1):194. doi:10.3390/ijms22010194

25. Wolswijk G. Oligodendrocyte precursor cells in the demyelinated multiple sclerosis spinal cord. Brain. 2002 Feb 1;125(2):338–49. doi:10.1093/brain/awf031

26. Paolicelli RC, Sierra A, Stevens B, Tremblay ME, Aguzzi A, Ajami B, et al. Microglia states and nomenclature: A field at its crossroads. Neuron. 2022 Nov;110(21):3458–83. doi:10.1016/j.neuron.2022.10.020

27. Reddaway J, Richardson PE, Bevan RJ, Stoneman J, Palombo M. Microglial morphometric analysis: so many options, so little consistency. Front Neuroinform. 2023 Aug 10;17:1211188. doi:10.3389/fninf.2023.1211188

28. Reddaway J, Richardson PE, Bevan RJ, Stoneman J, Palombo M. Microglial morphometric analysis: so many options, so little consistency. Front Neuroinform. 2023 Aug 10;17:1211188. doi:10.3389/fninf.2023.1211188

29. Green TRF, Rowe RK. Quantifying microglial morphology: an insight into function. Clinical and Experimental Immunology. 2024 May 16;216(3):221–9. doi:10.1093/cei/uxae023

30. Au NPB, Ma CHE. Recent Advances in the Study of Bipolar/Rod-Shaped Microglia and their Roles in Neurodegeneration. Front Aging Neurosci. 2017 May 4;9:128. doi:10.3389/fnagi.2017.00128

31. Yamada J, Jinno S. Novel objective classification of reactive microglia following hypoglossal axotomy using hierarchical cluster analysis. J of Comparative Neurology. 2013 Apr;521(5):1184–201. doi:10.1002/cne.23228

32. Lin HH, Faunce DE, Stacey M, Terajewicz A, Nakamura T, Zhang-Hoover J, et al. The macrophage F4/80 receptor is required for the induction of antigen-specific efferent regulatory T cells in peripheral tolerance. The Journal of Experimental Medicine. 2005 May 16;201(10):1615–25. doi:10.1084/jem.20042307

33. Walker DG, Lue LF. Immune phenotypes of microglia in human neurodegenerative disease: challenges to detecting microglial polarization in human brains. Alz Res Therapy. 2015 Dec;7(1):56. doi:10.1186/s13195-015-0139-9

34. Cantoni C, Bollman B, Licastro D, Xie M, Mikesell R, Schmidt R, et al. TREM2 regulates microglial cell activation in response to demyelination in vivo. Acta Neuropathol. 2015 Mar;129(3):429–47. doi:10.1007/s00401-015-1388-1

35. Poliani PL, Wang Y, Fontana E, Robinette ML, Yamanishi Y, Gilfillan S, et al. TREM2 sustains microglial expansion during aging and response to demyelination. J Clin Invest. 2015 May 1;125(5):2161–70. doi:10.1172/JCI77983

36. Wang Y, Kyauk RV, Shen YA, Xie L, Reichelt M, Lin H, et al. TREM2 -dependent microglial function is essential for remyelination and subsequent neuroprotection. Glia. 2023 May;71(5):1247–58. doi:10.1002/glia.24335

37. Gutiérrez IL, Martín-Hernández D, MacDowell KS, García-Bueno B, Caso JR, Leza JC, et al. CX3CL1 Regulation of Gliosis in Neuroinflammatory and Neuroprotective Processes. IJMS. 2025 Jan 23;26(3):959. doi:10.3390/ijms26030959

38. Xu S, Wang J, Zhong J, Shao M, Jiang J, Song J, et al. CD73 alleviates GSDMD-mediated microglia pyroptosis in spinal cord injury through PI3K/AKT/Foxo1 signaling. Clinical & Translational Med. 2021 Jan;11(1):e269. doi:10.1002/ctm2.269

39. Xu S, Zhu W, Shao M, Zhang F, Guo J, Xu H, et al. Ecto-5lll-nucleotidase (CD73) attenuates inflammation after spinal cord injury by promoting macrophages/microglia M2 polarization in mice. J Neuroinflammation. 2018 Dec;15(1):155. doi:10.1186/s12974-018-1183-8

40. Meng F, Guo Z, Hu Y, Mai W, Zhang Z, Zhang B, et al. CD73-derived adenosine controls inflammation and neurodegeneration by modulating dopamine signalling. Brain. 2019 Mar 1;142(3):700–18. doi:10.1093/brain/awy351

41. Shen K, Reichelt M, Kyauk RV, Ngu H, Shen YAA, Foreman O, et al. Multiple sclerosis risk gene Mertk is required for microglial activation and subsequent remyelination. Cell Reports. 2021 Mar;34(10):108835. doi:10.1016/j.celrep.2021.108835

42. Healy LM, Jang JH, Won SY, Lin YH, Touil H, Aljarallah S, et al. MerTK-mediated regulation of myelin phagocytosis by macrophages generated from patients with MS. Neurol Neuroimmunol Neuroinflamm. 2017 Nov;4(6):e402. doi:10.1212/NXI.0000000000000402

43. Burstyn-Cohen T, Fresia R. TAM receptors in phagocytosis: Beyond the mere internalization of particles. Immunological Reviews. 2023 Oct;319(1):7–26. doi:10.1111/imr.13267

44. Que X, Zhang T, Liu X, Yin Y, Xia X, Gong P, et al. The role of TREM2 in myelin sheath dynamics: A comprehensive perspective from physiology to pathology. Progress in Neurobiology. 2025 Apr;247:102732. doi:10.1016/j.pneurobio.2025.102732

45. Pocock J, Vasilopoulou F, Svensson E, Cosker K. Microglia and TREM2. Neuropharmacology. 2024 Oct;257:110020. doi:10.1016/j.neuropharm.2024.110020

46. Karadima E, Chavakis T, Alexaki VI. Arginine metabolism in myeloid cells in health and disease. Semin Immunopathol. 2025 Dec;47(1):11. doi:10.1007/s00281-025-01038-9

47. Sonar SA, Lal G. The iNOS Activity During an Immune Response Controls the CNS Pathology in Experimental Autoimmune Encephalomyelitis. Front Immunol. 2019 Apr 4;10:710. doi:10.3389/fimmu.2019.00710

48. Xu T, Liu C, Deng S, Gan L, Zhang Z, Yang GY, et al. The roles of microglia and astrocytes in myelin phagocytosis in the central nervous system. J Cereb Blood Flow Metab. 2023 Mar;43(3):325–40. doi:10.1177/0271678X221137762

49. Cunniffe N, Coles A. Promoting remyelination in multiple sclerosis. J Neurol. 2021 Jan;268(1):30–44. doi:10.1007/s00415-019-09421-x

50. Baaklini CS, Ho MFS, Lange T, Hammond BP, Panda SP, Zirngibl M, et al. Microglia promote remyelination independent of their role in clearing myelin debris. Cell Reports. 2023 Dec;42(12):113574. doi:10.1016/j.celrep.2023.113574

51. Wagner J, Hoyer C, Antony H, Lundgrén K, Soliymani R, Crux S, et al. CX3CR1 modulates migration of resident microglia towards brain injury [Internet]. Neuroscience; 2024 [cited 2026 Feb 10]. Available from: http://biorxiv.org/lookup/doi/10.1101/2024.09.23.614458 doi:10.1101/2024.09.23.614458

52. Rivest S. CX3CR1 in multiple sclerosis. Oncotarget. 2015 Aug 21;6(24):19946–7. doi:10.18632/oncotarget.4650

53. Ong SM, Hadadi E, Dang TM, Yeap WH, Tan CTY, Ng TP, et al. The pro-inflammatory phenotype of the human non-classical monocyte subset is attributed to senescence. Cell Death Dis. 2018 Feb 15;9(3):266. doi:10.1038/s41419-018-0327-1

54. Smith JA, Das A, Ray SK, Banik NL. Role of pro-inflammatory cytokines released from microglia in neurodegenerative diseases. Brain Res Bull. 2012 Jan 4;87(1):10–20. doi:10.1016/j.brainresbull.2011.10.004 PubMed PMID: 22024597; PubMed Central PMCID: PMC9827422.

55. Cignarella F, Filipello F, Bollman B, Cantoni C, Locca A, Mikesell R, et al. TREM2 activation on microglia promotes myelin debris clearance and remyelination in a model of multiple sclerosis. Acta Neuropathol. 2020 Oct;140(4):513–34. doi:10.1007/s00401-020-02193-z

56. Gouna G, Klose C, Bosch-Queralt M, Liu L, Gokce O, Schifferer M, et al. TREM2-dependent lipid droplet biogenesis in phagocytes is required for remyelination. Journal of Experimental Medicine. 2021 Oct 4;218(10):e20210227. doi:10.1084/jem.20210227

57. Hou J, Magliozzi R, Chen Y, Wu J, Wulf J, Strout G, et al. Acute TREM2 inhibition depletes MAFB-high microglia and hinders remyelination. Proc Natl Acad Sci USA. 2025 Apr;122(13):e2426786122. doi:10.1073/pnas.2426786122

58. Morell P, Quarles RH. Chapter 4: Myelin Formation, Structure and Biochemistry. In: Basic Neurochemistry: Molecular, Cellular and Medical Aspects. 6th edition. [Internet]. 6th edition. Philadelphia, Pennsylvania, USA: Lippincott-Raven Publishers; 1999. Available from: https://www.ncbi.nlm.nih.gov/books/NBK20402/

59. Mather ML, Evangelou AV, Bourne JN, Macklin WB, Wood TL. Myelin Lipid Composition in the Central Nervous System Is Regionally Distinct and Requires Mechanistic Target of Rapamycin Signaling. Glia. 2025 Sep;73(9):1841–59. doi:10.1002/glia.70042 PubMed PMID: 40417825; PubMed Central PMCID: PMC12313007.

60. Marschallinger J, Iram T, Zardeneta M, Lee SE, Lehallier B, Haney MS, et al. Lipid-droplet-accumulating microglia represent a dysfunctional and proinflammatory state in the aging brain. Nat Neurosci. 2020 Feb;23(2):194–208. doi:10.1038/s41593-019-0566-1 PubMed PMID: 31959936; PubMed Central PMCID: PMC7595134.

61. Prakash P, Manchanda P, Paouri E, Bisht K, Sharma K, Rajpoot J, et al. Amyloid-β induces lipid droplet-mediated microglial dysfunction via the enzyme DGAT2 in Alzheimer’s disease. Immunity. 2025 Jun;58(6):1536-1552.e8. doi:10.1016/j.immuni.2025.04.029

62. Mahmood A, Miron VE. Microglia as therapeutic targets for central nervous system remyelination. Current Opinion in Pharmacology. 2022 Apr;63:102188. doi:10.1016/j.coph.2022.102188

63. Zhong L, Chen XF, Wang T, Wang Z, Liao C, Wang Z, et al. Soluble TREM2 induces inflammatory responses and enhances microglial survival. Journal of Experimental Medicine. 2017 Mar 6;214(3):597–607. doi:10.1084/jem.20160844

64. Cai B, Thorp EB, Doran AC, Subramanian M, Sansbury BE, Lin CS, et al. MerTK cleavage limits proresolving mediator biosynthesis and exacerbates tissue inflammation. Proc Natl Acad Sci USA. 2016 Jun 7;113(23):6526–31. doi:10.1073/pnas.1524292113

